# *Crocosphaera* as a major consumer of fixed nitrogen despite its capability of nitrogen fixation

**DOI:** 10.1101/2021.07.28.454264

**Authors:** Takako Masuda, Keisuke Inomura, Taketoshi Kodama, Takuhei Shiozaki, Satoshi Kitajima, Gabrielle Armin, Takato Matsui, Koji Suzuki, Shigenobu Takeda, Ondřej Prášil, Ken Furuya

## Abstract

*Crocosphaera watsonii* (hereafter *Crocosphaera*) is a key nitrogen (N) fixer in the ocean, but its ability to consume combined N sources is still unclear. Using *in situ* microcosm incubations with an ecological model, we show that *Crocosphaera* has high competitive capability both under low and moderately high combined N concentrations. In field incubations, *Crocosphaera* accounted for the highest consumption of ammonium and nitrate, followed by pico-eukaryotes. The model analysis shows that cells have a high ammonium uptake rate (∼7 mol N (mol N)^-1^ d^-1^ at the maximum), which allows them to compete against pico-eukaryotes and non-diazotrophic cyanobacteria when combined N is sufficiently available. Even when combined N is depleted, their capability of nitrogen fixation allows higher growth rates compared to potential competitors. These results suggest the high fitness of *Crocosphaera* in combined N limiting, oligotrophic oceans, and thus heightens its potential significance in its ecosystem and in biogeochemical cycling.

## Introduction

Marine phytoplankton contribute about one half of the global net primary production and play a key role in regulating global biogeochemical cycles (1). Since phytoplankton are biochemically, metabolically, and ecologically diverse (2-4), understanding the contribution of different phytoplankton groups to ecosystem functioning is central to the precise estimation of the global carbon (C) and nitrogen (N) budget and in predicting the biogeochemical impact of future environmental changes (5).

In the oligotrophic subtropical gyres, combined N (defined as N covalently bonded to one or more elements other than N (6)) limits primary production and controls planktonic community composition (7-10). Therefore, N_2_ fixing microorganisms (diazotrophs) are important as a source of combined N in oligotrophic ecosystems (11, 12). In the subtropic oligotrophic ocean, the unicellular diazotroph, *Crocosphaera watsonii* (2.5 – 6 µm), is widely distributed (10, 13-16) in addition to pico-sized (<3 µm) cyanobacteria (e.g., *Prochlorococcus* and *Synechococcus*) and pico-eukaryotes (17-19). Recent studies reveal *Crocosphaera watsonii*’s ability to assimilate dissolved inorganic nitrogen (DIN), such as ammonium (NH_4_^+^) and nitrate (NO_3_^-^), at a nanomolar level and keep fixing N_2_ (20, 21). Model results indicate using DIN enables *Crocosphaera* to increase their abundance and expand their niche (22). These studies proposed that unicellular diazotrophs can be competitors with non-diazotrophic phytoplankton for combined N. However, how *Crocosphaera* competes for combined N is poorly evaluated. In this study, we combine an *in situ* microcosm experiment with N addition at the nanomolar level and model (23) to evaluate the competitiveness of *Crocosphaera* in a N limiting environment.

## Results

### Summary of the experiment

We carried out five nitrogen (N) and phosphorus (P)-addition bioassays (M1 to M5) at a station in the subtropical Northwestern Pacific (12°N, 135°E) from 6 to 25 June 2008 during the MR08-02 cruise on the R/V *MIRAI*. Nutrient concentrations initially were less than 36 nM for ammonium (NH_4_^+^), 7 nM for nitrate plus nitrite (NO_3_^-^ + NO_2_^-^) and 64 nM for phosphorus (PO_4_^3-^) (24). The physical and biological parameters at the initial condition of the experiments are described in (24). Hydrography and biochemistry at the station are described in (25). Although we performed pre-filtration with a 1 µm-filter to eliminate the effect of grazing, water samples contained plankton with up to ∼5 µm in size.

### Nutrient uptake and fate of enriched DIN

For 3 days of incubation, the phytoplankton community consumed NH_4_^+^ entirely at the end, while NO_3_^-^ was not always consumed completely (Fig. 1, Fig. S1). Estimated biomass explains about half of consumed combined N sources (Figs. 1, 2A), possibly due to luxury uptake (26, 27).

**Fig. 1.**
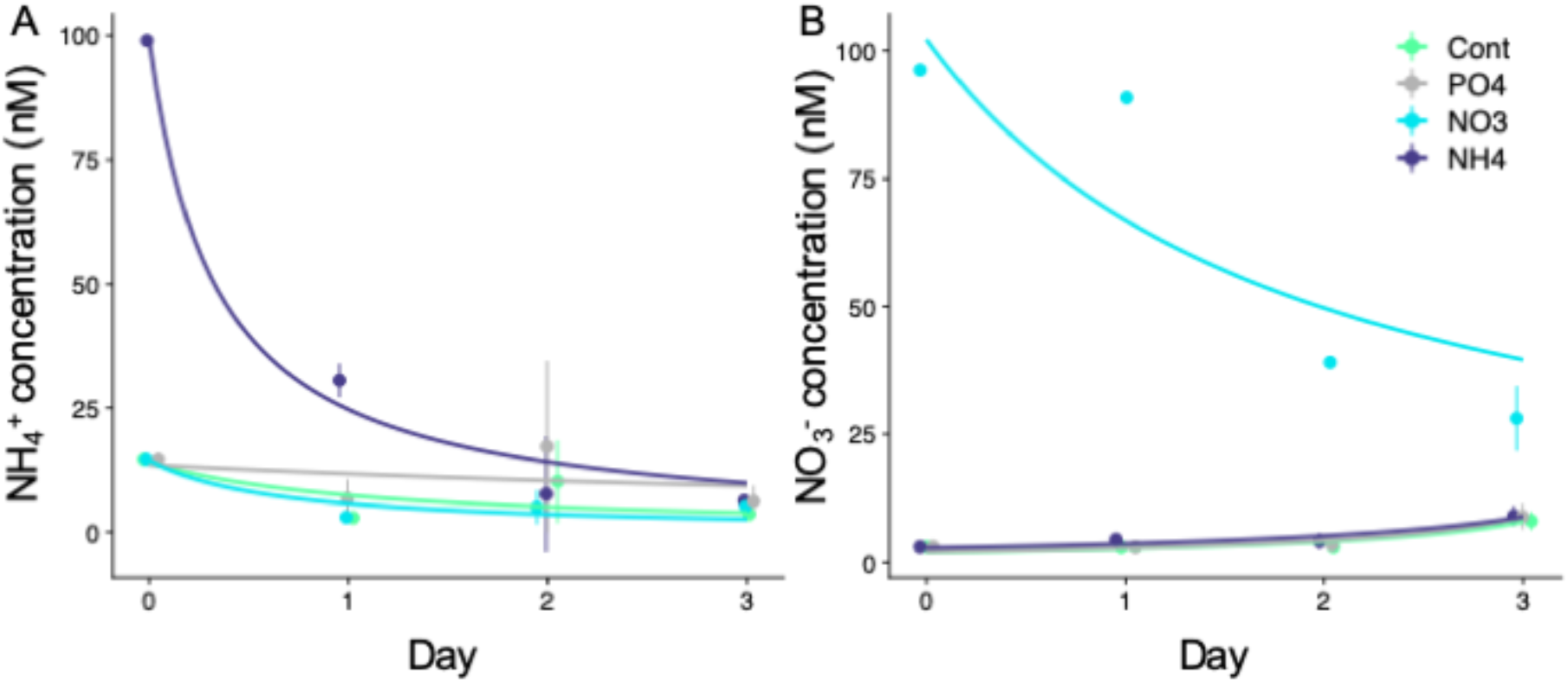
Temporal change in NH_4_^+^ and NO_3_^-^ concentrations of Ex. M3. (A) NH_4_^+^ concentration in the NH_4_^+^ treatment exponentially decreased during the experiment down to the detection limit of 6 nM on day 3. (B) NO_3_^-^ concentrations in the NO_3_^-^ treatment exponentially decreased during the experiment, but enriched NO_3_^-^ was not always entirely consumed. Error bar shows a standard deviation of triplicate. Temporal change in Urea-N concentration is shown in Fig. S2.

**Fig. 2.**
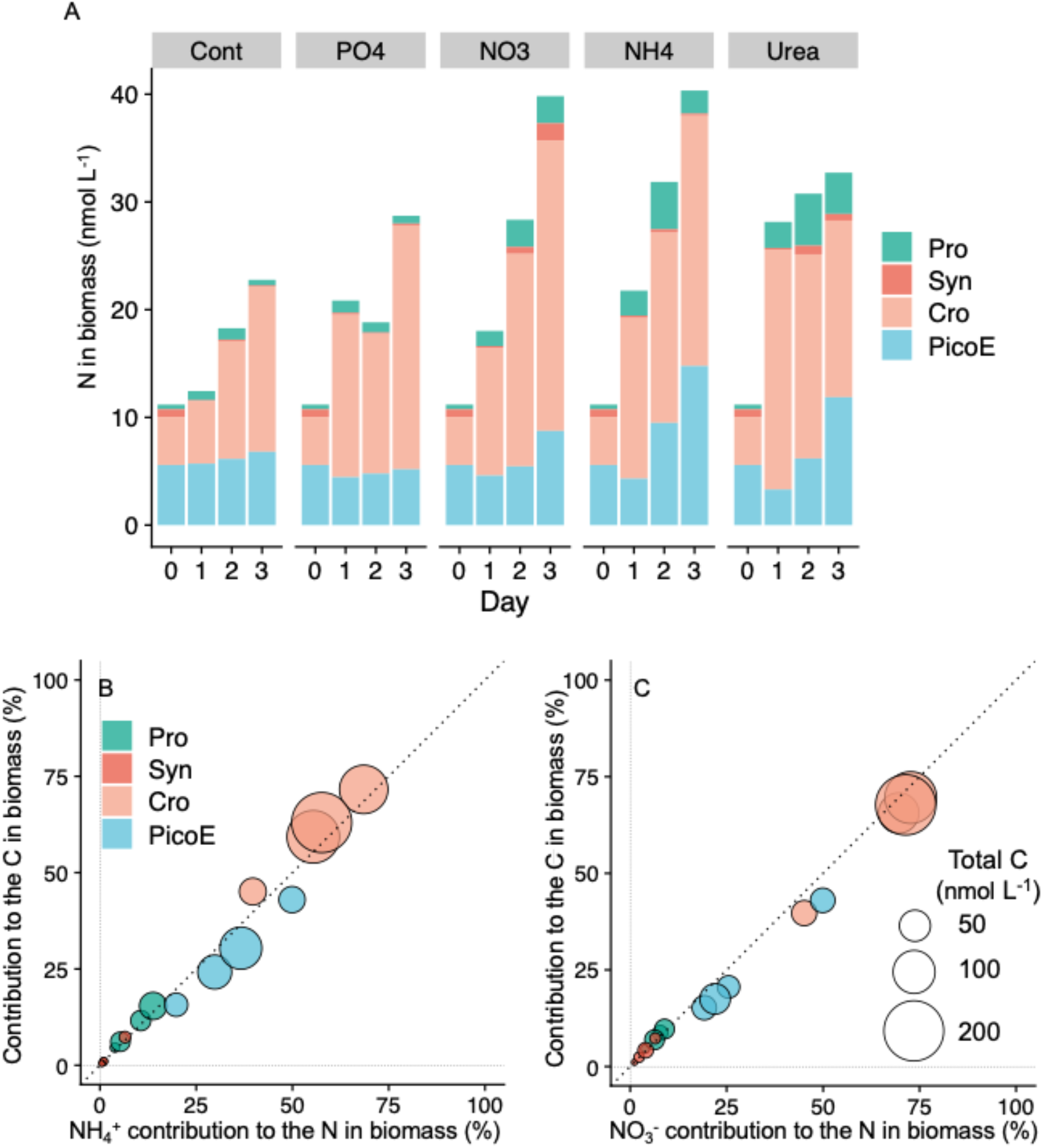
(A) N in biomass in each treatment and its contribution of each phytoplankton group of experiment M3. (B) Contribution to total carbon C in biomass as a function of the contribution of NH_4_^+^ -N biomass for each phytoplankton group. (C) Contribution to total carbon C in biomass as a function of the contribution of NO_3_^-^ -N biomass for each phytoplankton group. Each circle shows data from a different day, and the size of the dots represents the total C in biomass (nmol C L^-1^). Pro; *Prochlorococcus*, Syn; *Synechococcus*, Cro; *Crocosphaera*, PicoE; pico-eukaryotes.

The greatest portion of estimated C and N in biomass were found in *Crocosphaera* (39-93% in all N addition incubations) followed by pico-eukaryotes (5-55% in N addition incubations) (Fig. 2A, Fig. S3). Although the origin of water mass changed from oligotrophic-water to mixed-water between experiments (Exs.) M1-M3 and M4-M5 (25), with more *Crocosphaera* in cell density at the latter environment (Table S1), the dominance of *Crocosphaera* as a C and N biomass was observed from all the experiments. N derived from N_2_ fixation was not always sufficient to support the N demand of *Crocosphaera*, especially in N amendment (Fig. S4). Estimated N_2_ fixation supported 0.5 – 12.7% of N demand of *Crocosphaera* in control and 0.5 – 11.6% in NH_4_^+^ treatment (Fig. S4), suggesting that *Crocosphaera* consumed amended N sources. Assimilation of combined nitrogen (NH_4_^+^ and NO_3_^-^), together with N fixation by *Crocosphaera*, has been reported earlier (20, 21). Although enriched 100 nM NH_4_^+^ was completely consumed (< 6 nM; detection limit, on day 3), increases in N-biomass of non-diazotrophs for 3 days were limited to up to 58 nmol L^-1^, again suggesting *Crocosphaera* took up combined nitrogen.

### Model analysis of the data

To quantitatively interpret the observed data, we used a simple model of the cellular growth, which is based on the uptake of NH_4_^+^ and NO_3_^-^ (see Methods).

We used the data from experiment M3 since it shows the clearest trends with low initial nutrient concentrations. The model captured the overall trend of the transition of cellular N (Fig. 3) based on the available nutrient (Fig. S5). The parameterization of the model reveals high rates of N uptake by *Crocosphaera*. Especially, we used about 7 (mol N (mol N)^-1^ d^-1^) for maximum NH_4_^+^ uptake to represent the data, which shows high combined-N uptake compared to other phytoplankton. Specifically, such parameterization was needed to reproduce the rapid growth of *Crocosphaera* under NH_4_^+^ added case between day 0 and day 1. The predicted maximum NO_3_^-^ uptake rate for *Crocosphaera* is also higher than for other phytoplankton, which is supported by *Crocosphaera*’s faster growth with NO_3_^-^ addition.

**Fig. 3.**
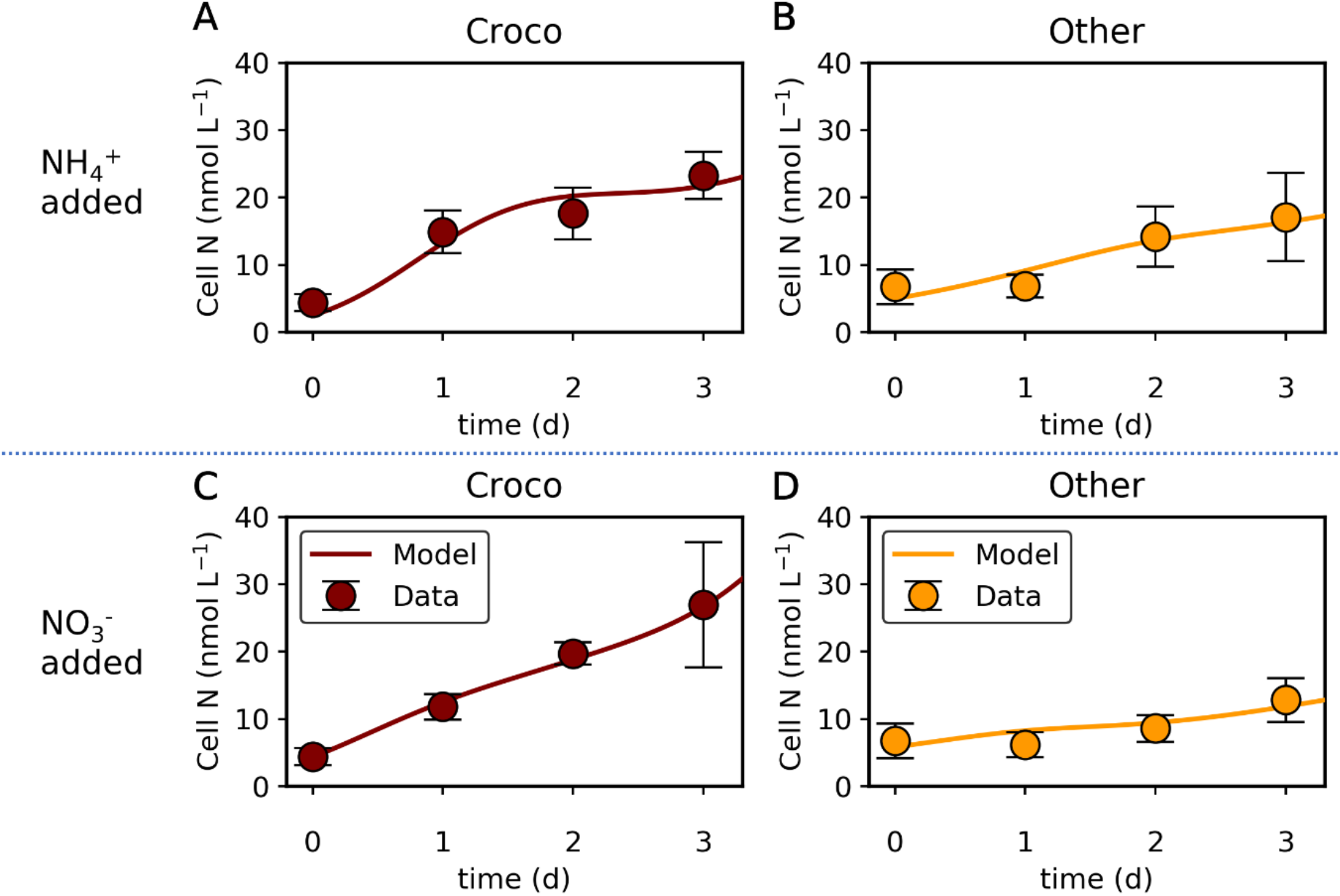
Simulated transition of cellular N with nutrient addition compared with data. (A)(B) NH_4_^+^ added case. (C)(D) NO_3_^-^ added case. Croco: *Crocosphaera*. Other: other phytoplankton. Data are from experiment M3.

To test the competitiveness of *Crocosphaera*, we simulated a simple ecological situation. Here, we simulate zooplankton with Kill the Winner Theory (KTW) (28), which is based on a commonly observed active prey-switching behavior of zooplankton (29-31). The result shows the high competitiveness of *Crocosphaera* both under high and low nutrient concentrations. Under high nutrient concentration, *Crocosphaera* may dominate other phytoplankton due to the high rate of nutrient uptake (Fig. 4A, S6B). However, under extremely low nutrient conditions (NH_4_^+^ and NO_3_^-^ are both at 1 nmol L^-1^), *Crocosphaera* is slightly outcompeted (Fig. 4B, S6B). This is due to the relatively high half-saturation constant for NH_4_^+^, which is manifested by the sudden decrease in growth rate with a drop in NH_4_^+^ under NH_4_^+^ addition (Fig. 3A, S5A). However, this relationship flips if we consider the effect of N_2_ fixation, which maintains their growth rates at a higher level rather than relying on external N under N depletion (Fig. 4C, S6C). These results suggest that possession of nitrogenase (an enzyme complex involved in N_2_ fixation) allows for *Crocosphaera’s* survival under low nutrient environments.

**Fig. 4.**
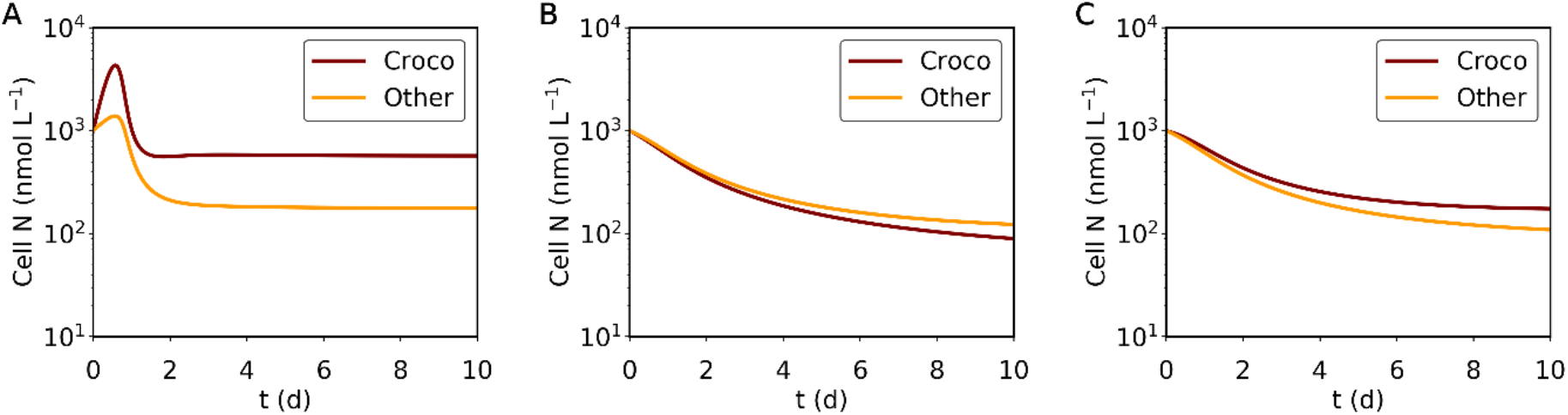
Simulated transition of cellular N in a simple ecosystem model for three different scenarios. (A) The concentrations for NH_4_^+^ and NO_3_^-^ are both 100 nmol L^-1^ (B)(C) The concentrations for NH_4_^+^ and NO_3_^-^ are both 1 nmol L^-1^. In only (C) *Crocosphaera* may acquire N via N_2_ fixation; in (A) and (B) the effect of N_2_ fixation is neglected. Croco: *Crocosphaera*. Other: other phytoplankton. Parameters are based on NH_4_^+^ added case.

## Discussion

Our study shows high uptake of N by *Crocosphaera* under relatively high N concentration. The results counter the general image of *Crocosphaera* since it is mostly known as a diazotroph and is considered to be a provider of N to the environment. Rather, our result supports more recent studies, where *Crocosphaera* does not increase the productivity of other phytoplankton (32) or even compete with other species over combined N (22). Surprisingly, our study even shows higher maximum uptake rates of NH_4_^+^ and NO_3_^-^, which allow its dominance just by uptake of combined N. When nitrogen concentration is extremely low, they could be outcompeted in N uptake, but their N_2_ fixation allows maintaining *Crocosphaera* biomass at a certain level, which can still be higher than those of non-diazotrophic phytoplankton. This high consumption of NO_3_^-^ may differ from UCYN-A (15, 33-35), which keeps fixing nitrogen under high NO_3_^-^ availability (36, 37), leading to their unique niche acquisition. These results suggest that *Crocosphaera* has high competitiveness under both low and high nutrients.

Despite that, we generally do not observe the oligotrophic ocean completely dominated by *Crocosphaera*. One reason might be the grazing selection. *Crocosphaera* is a unicellular cyanobacterium a few microns to 6 µm in diameter (38), and its tight coupling with predators is reported recently (39). The new production of *Crocosphaera* is estimated to support up to 400% of C demand of the main grazers, and the grazing rates of the main predator *Protoperidinium* were found to be nearly equivalent to growth rates of *Crocosphaera* (39). On the other hand, its potential competitor, *Trichodesmium*, a major N_2_ fixer in the ocean, is reported to produce a toxin (40-42), and creates large colonies of ∼10^4^ cells (43), potentially protecting themselves from grazing. Another reason might be the growth limitation by other nutrients such as P and Fe. Although there are some reports that *Crocosphaera* shows adaptation for low P and low Fe, their relative fitness to such low P or low Fe environments compared to other organisms has not been quantified. Since having nitrogenase enzymes require a high concentration of Fe, non-nitrogen fixers, such as *Prochlorococcus* and *Synechococcus*, may have lower Fe requirements and are more adapted to Fe depletion. Also, *Crocosphaera* does not seem to fully utilize sulpholipid, which would save P use, as opposed to other cyanobacteria, such as *Synechococcus* (44, 45), and thus may not compete strongly under P limitation.

At the same time, it is largely possible that *Crocosphaera* dominates at some regions in the oligotrophic ocean given its high competitiveness under N limitation, which is the characteristic of the oligotrophic ocean (7, 46). For example, a study of flow cytometry shows a high abundance of *Crocosphaera*-like cells in a wide region of the North Pacific (47), where the abundance of *Trichodesmium* seems limited (48). Also, a recent study shows multiple gene copies of *Trichodesmium* (up to ∼700 gene copied per cell) (49), which would overestimate their abundance (50). Given these factors and our analysis showing their high fitness to both low and high nitrogen concentration, it is possible that we are still underestimating the relative abundance and role of *Crocosphaera* in global biogeochemical cycling.

## Materials and methods

### Experimental setup and sample collection

The dataset presented herein originates from an experimental setup described in (24). Briefly, we carried out five macro-nutrient (N and P)-addition bioassays (M1 to M5) using natural phytoplankton assemblages collected at a station in the subtropical Northwestern Pacific (12°N, 135°E) from 6-25 June 2008 during the MR08-02 cruise on the R/V *MIRAI*. For macro-nutrient bioassays, we distributed pre-filtered seawater from 10 m depth into 4-liter polycarbonate bottles. We performed three treatments with 100 nM addition of N as NaNO_3_, NH_4_Cl, or urea, and one treatment with 10 nM of NaH_2_PO_4_. Our control was without nutrient addition. Bottles were incubated on deck for three days with daily sample harvest in flow-through seawater tanks covered with a neutral density screen to attenuate light intensity to 50% of its corresponding surface value.

### Macro-nutrient and iron concentrations

Concentrations of NO_3_^-^ +NO_2_^-^(N + N), NH_4_^+^, Soluble Reactive Phosphorus (SRP), and urea were measured using a high-sensitivity colorimetric approach with an AutoAnalyzer II (Technicon) and Liquid Waveguide Capillary Cells (World Precision Instruments, USA) as outlined (51). We analyzed urea concentrations using the diacetyl monoxime method (52). Detection limits of NO_3_^-^ + NO_2_^-^, NH_4_^+^, and SRP were 3, 6, and 3 nM, respectively.

### Flow cytometry

Flow-cytometry (FCM) identified *Prochlorococcus, Synechococcus*, pico-eukaryotes, and *Crocosphaera* based on cell size and chlorophyll- or phycoerythrin-fluorescence. Aliquots of 4.5 mL were preserved in glutaraldehyde (1% final concentration), flash-frozen in liquid N_2_, and stored at -80 °C until analysis on land by flow cytometry (PAS-III, Partec, GmbH, Münster, Germany) equipped with a 488 nm argon-ion excitation laser (100 mW). We recorded forward- and side-angle scatter (FSC and SSC), red fluorescence (>630 nm, FL3), and orange fluorescence (570–610 nm, FL2). FloMax® (Partec, GmbH, Münster, Germany) distinguished *Synechococcus, Prochlorococcus, Crocosphaera*, and pico-eukaryotes based on their auto-fluorescence properties and their size.

### Gene analysis

We collected DNA samples from each treatment of the Fe addition bioassay and collected aliquots of 0.5 to 1.0 L of sample on 0.2 µm SUPOR® polyethersulfone membrane filters, which we then placed in sterile tubes containing glass beads, frozen in liquid N_2_, and stored at -80°C until further analysis. DNA was extracted according to (53) to determine the abundance of *Crocosphaera watsonii* by quantitative PCR (qPCR) using a 5’ nuclease assay as described in (54).

Quantitative PCR showed that cell densities of FCM-identified *Crocosphaera* were significantly, positively correlated with *nif*H gene copies used to quantify the proportion of *Crocosphaera*, indicating that *nifH* abundance accounted for 68% of the variation in FCM-identified *Crocosphaera* (r^2^ = 0.463, n = 48, *p*=0.001, Pearson Product Moment correlation). Therefore, this study treated FCM-identified *Crocosphaera* as diazotroph *Crocosphaera*. Cell abundance estimated by qPCR was 0.63 ± 0.23 fold lower than those measured by FCM.

### Nitrogen fixation

To measure *in situ* N_2_ fixation activity, we used the acetylene reduction assay of (55, 56). We dispensed a total of 550 milliliter bioassay samples into 1200 mL HCl– rinsed glass PETG bottles with 6 replicates and sealed with butyl rubber stoppers. Aliquots of 120 mL of acetylene (99.9999% (v:v), Kouatsu Gas Kogyo, Japan) were injected through the stopper by replacing the same volume of headspace. After 24 h in the on-deck flow-through seawater tanks, we analyzed ethylene concentrations by converting the ethylene to fixed nitrogen with a molar ratio of 4:1 (57).

### Cellular C and N estimation

We used a conversion factor of 235 fg Cµm^-3^ for *Prochlorococcus, Synechococcus* and *Crocosphaera* (58) to estimate cellular carbon content. For picoeukaryotes, we represented cell volume by converting it into carbon per cell, using a modified Strathmann equation (58, 59):

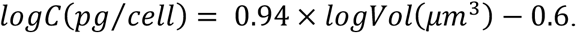

Then, using an earlier reported C:N ratio (C:N ratio = 9.1 for *Prochlorococcus*, 8.6 for *Synechococcus*, 8.7 for *Crocosphaera*, 6.6 for picoeukaryotes), we converted the cellular C content into cellular N (21, 60, 61).

### Statistical analysis

Phytoplankton cell densities of each bioassay were first compared between treatments using repeated measurements Analysis of Variance (RM-ANOVA) with nutrient treatments as a between-subjects factor (5 levels) and time (4 levels) as the within-subjects factor. Treatment effects were considered significant if *p* < 0.05. Then, means between five treatments were compared by post hoc Turkey test (n = 3 replicates per treatment throughout, degrees of freedom = 40).

### Quantitative model of microbial growth

To quantitatively analyze the fitness of *Crocosphaera* under N limiting conditions, we ran two simulations. One was to represent the incubation experiment to extract parameters manually and the other was the simple ecosystem model to simulate their competitiveness under different nutrient concentrations and scenarios. The list of parameters and used values are in Table S2 and S3, respectively.

#### Simulation of the incubation experiment

We used the following equations for the growth of phytoplankton to represent the field incubation experiment:

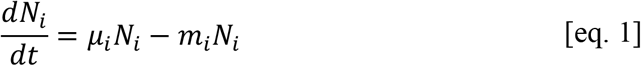

where *N*_*i*_ (nmol L^-1^) is the cellular nitrogen concentration of phytoplankton *i* (i = Cro, Oth: *Crocosphaera* and other phytoplankton, respectively) per volume water, *t* (d) is time, *µ*_*i*_ (d^-1^) is the growth rate of phytoplankton i, and *m*_*i*_ (d^-1^) is a mortality rate of phytoplankton i.

To represent the growth of *Crocosphaera* and other phytoplankton, we used simple growth equations based on the sum of Monod kinetics (62) for each nutrient:

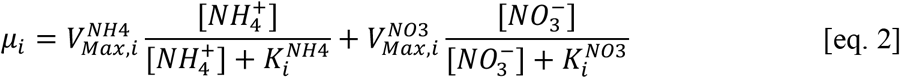

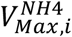 and 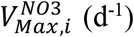 are the maximum uptake rate of phytoplankton for NH_4_^+^ and NO_3_^-^ respectively, [*j*] (nmol L^-1^) is the concentration of nutrient 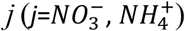, and 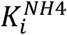 and 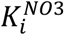 (nmol L^-1^) are half-saturation constant of nutrient for phytoplankton i respectively. We used the data-fitted quadratic curve of nutrient concentrations (Fig. S5).

#### Simple ecosystem simulation

To simulate the simple ecosystem situation, we introduced the grazing by zooplankton:

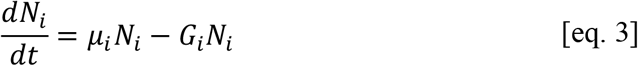

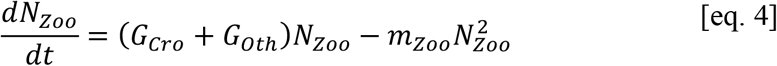

where *G*_i_ (d^-1^) is the grazing rate of phytoplankton *i* by zooplankton, *N*_*Zoo*_ (nmol L^-1^) is the nitrogen concentration in zooplankton per volume water, and *M*_Z*oo*_ (d^-2^) is a quadratic mortality rate of zooplankton. When we allow nitrogen fixation, we used *µ*_Cro_ = 0.31 (d^-1^) (a typical growth rate under diazotrophic conditions (63), if the computation based on [eq.2] yields a value below 0.31 (d^-1^).

For *G*_i_ we have applied the KTW method:

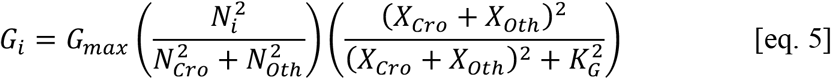

where *G*_m*ax*_ (d^-1^) is the maximum grazing rate and *K*_Y_ (nmol L^-1^) is grazing half-saturation. This equation reflects the commonly observed prey-switching behavior of zooplankton (29-31), which stabilizes ecosystems (64, 65).

## Code availability

The model developed in this paper has been uploaded in GitHub/Zenodo and is freely available at https://zenodo.org/record/5095790 (DOI: 10.5281/zenodo.5095790).

## Author contributions

T.Masuda, K.F and S.T designed the *in situ* microcosm experiments, T.Masuda. T.K, T.S, S.K, and T.Matsui carried out the experiment and analyzed data supervised by K.F, S.T and K.S. T.Masuda and K.I shaped the concept of the study with the supervision of O.P. K.I. and G.A. developed and ran the model. T.Masuda and K.I. wrote the original draft with substantial input from all the authors.

## Acknowledgements

We thank the captain, crew and technicians of the R/V *MIRAI* for assistance and support during the research cruise. This research was financially supported by MEXT grants for Scientific Research on Innovative Areas (24121001, 24121005, K.F.), Czech Research Foundation GAČR (project 20-17627S to O.P. and T.Masuda), the Simons Foundation (Life Sciences-Simons Postdoctoral Fellowships in Marine Microbial Ecology, Award 544338, K.I.), and the National Science Foundation (NSF) under EPSCoR Cooperative Agreement (#OIA-1655221, K.I.).

---

**Table S1.** *In situ* nitrogen fixation rate at 10 m depth and cell density of *Crocosphaera* in the incubation bottle at initial. For all data, means are shown with ± standard deviation for triplicate samples.

| Ex. | Date | *In situ* N<sub>2</sub> fixation rate (nmolN L<sup>-1</sup>d<sup>-1</sup>) | *In situ* N<sub>2</sub> fixation rate <10μm (nmolN L<sup>-1</sup>d<sup>-1</sup>) | *Crocospaera* at initial (cells mL<sup>-1</sup>) |
| --- | --- | --- | --- | --- |
|  | In 2008 |  |  |  |
| M1 | 6 June | 1.33 ± 1.81 | 2.75 ± 4.68 | 32 ± 62 |
| M2 | 10 June | 2.37 ± 0.59 | 0.66 ± 0.96 | 270 ± 225 |
| M3 | 14 June | 0.19 ± 2.06 | 0.28 ± 2.40 | 126 ± 32 |
| M4 | 18 June | 6.65 ± 2.52 | 2.37 ± 0.77 | 1513 ± 684 |
| M5 | 22 June | 4.94 ± 1.38 | 4.75 ± 1.72 | 306 ± 112 |

**Table S2.**
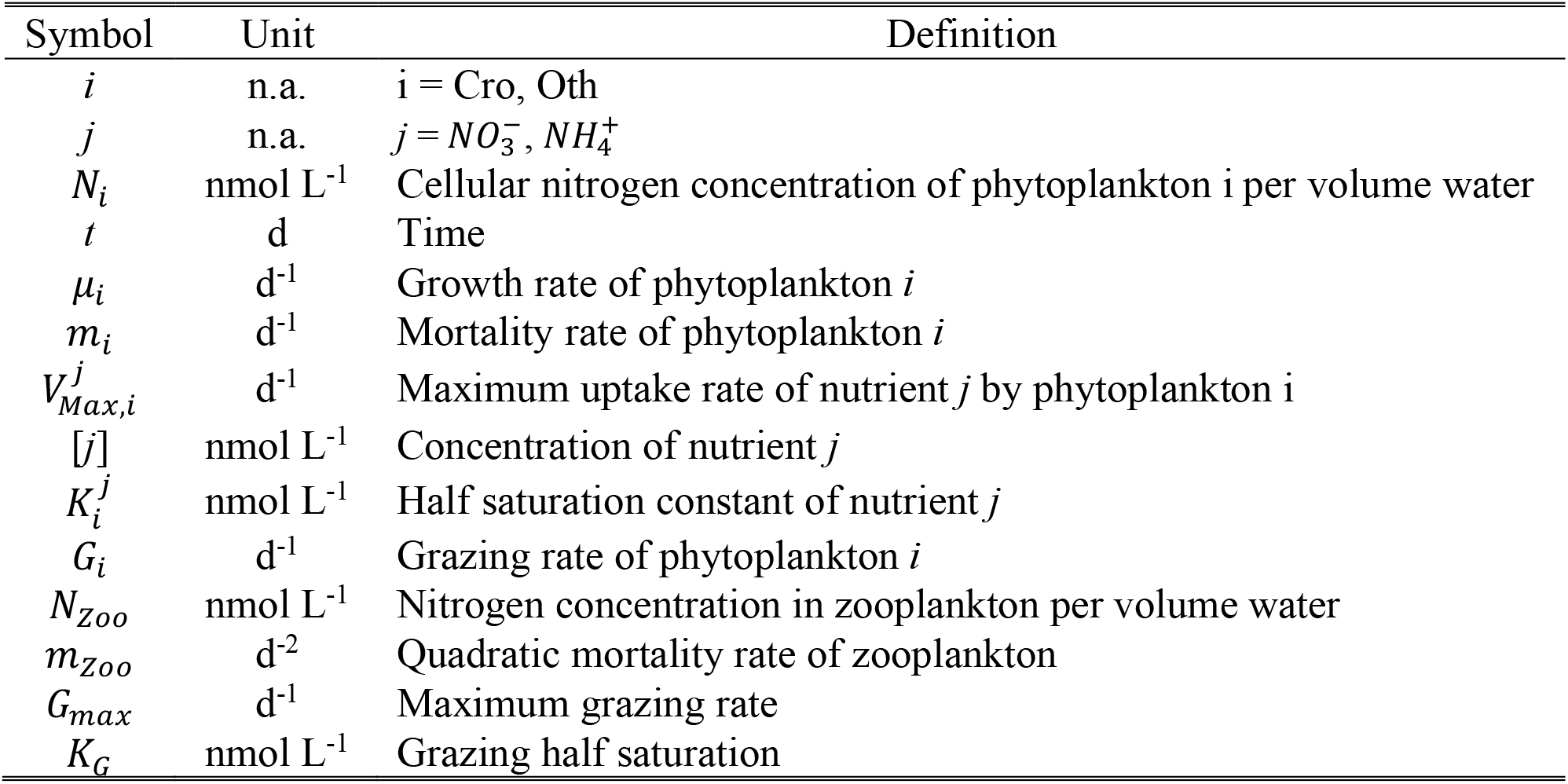
Used symboles, units and definitions in the quantitative model

**Table S3.** Values used for parameters

| Parameter | Unit | Value |
| --- | --- | --- |
| $m_i$ | $d^{-1}$ | 0.4 |
| For $NH_4^+$ added case | | |
| $N_{Cro}$ | $nmol\ L^{-1}$ | 2.50* |
| $N_{Oth}$ | $nmol\ L^{-1}$ | 5.00* |
| $V_{Max,Cro}^{NH_4}$ | $d^{-1}$ | 6.6 |
| $V_{Max,Cro}^{NO_3}$ | $d^{-1}$ | 2.8 |
| $V_{Max,Oth}^{NH_4}$ | $d^{-1}$ | 1.1 |
| $V_{Max,Oth}^{NO_3}$ | $d^{-1}$ | 1.8 |
| $K_{Cro}^{NH_4}$ | $nmol\ L^{-1}$ | 140 |
| $K_{Cro}^{NO_3}$ | $nmol\ L^{-1}$ | 80 |
| $K_{Oth}^{NH_4}$ | $nmol\ L^{-1}$ | 6 |
| $K_{Oth}^{NO_3}$ | $nmol\ L^{-1}$ | 500 |
| For $NO_3^-$ added case | | |
| $N_{Cro}$ | $nmol\ L^{-1}$ | 4.36* |
| $N_{Oth}$ | $nmol\ L^{-1}$ | 5.83*s |
| $V_{Max,Cro}^{NH_4}$ | $d^{-1}$ | 8 |
| $V_{Max,Cro}^{NO_3}$ | $d^{-1}$ | 1.3 |
| $V_{Max,Oth}^{NH_4}$ | $d^{-1}$ | 0.9 |
| $V_{Max,Oth}^{NO_3}$ | $d^{-1}$ | 0.5 |
| $K_{Cro}^{NH_4}$ | $nmol\ L^{-1}$ | 70 |
| $K_{Cro}^{NO_3}$ | $nmol\ L^{-1}$ | 90 |
| $K_{Oth}^{NH_4}$ | $nmol\ L^{-1}$ | 2 |
| $K_{Oth}^{NO_3}$ | $nmol\ L^{-1}$ | 700 |
| Ecosystem simulation |  |  |
| $N_{Cro}$ | $nmol\ L^{-1}$ | 1000* |
| $N_{Oth}$ | $nmol\ L^{-1}$ | 1000* |
| $N_{Zoo}$ | $nmol\ L^{-1}$ | 1000* |
| $m_{Zoo}$ | $d^{-2}$ | 0.01<sup>#</sup> |
| $G_{max}$ | $d^{-1}$ | 7.5 |
| $K_G$ | $d^{-1}$ | 500<sup>#</sup> |

\* Initial value. <sup>#</sup>Value from Inomura et al (2019).

**Fig. S1.**
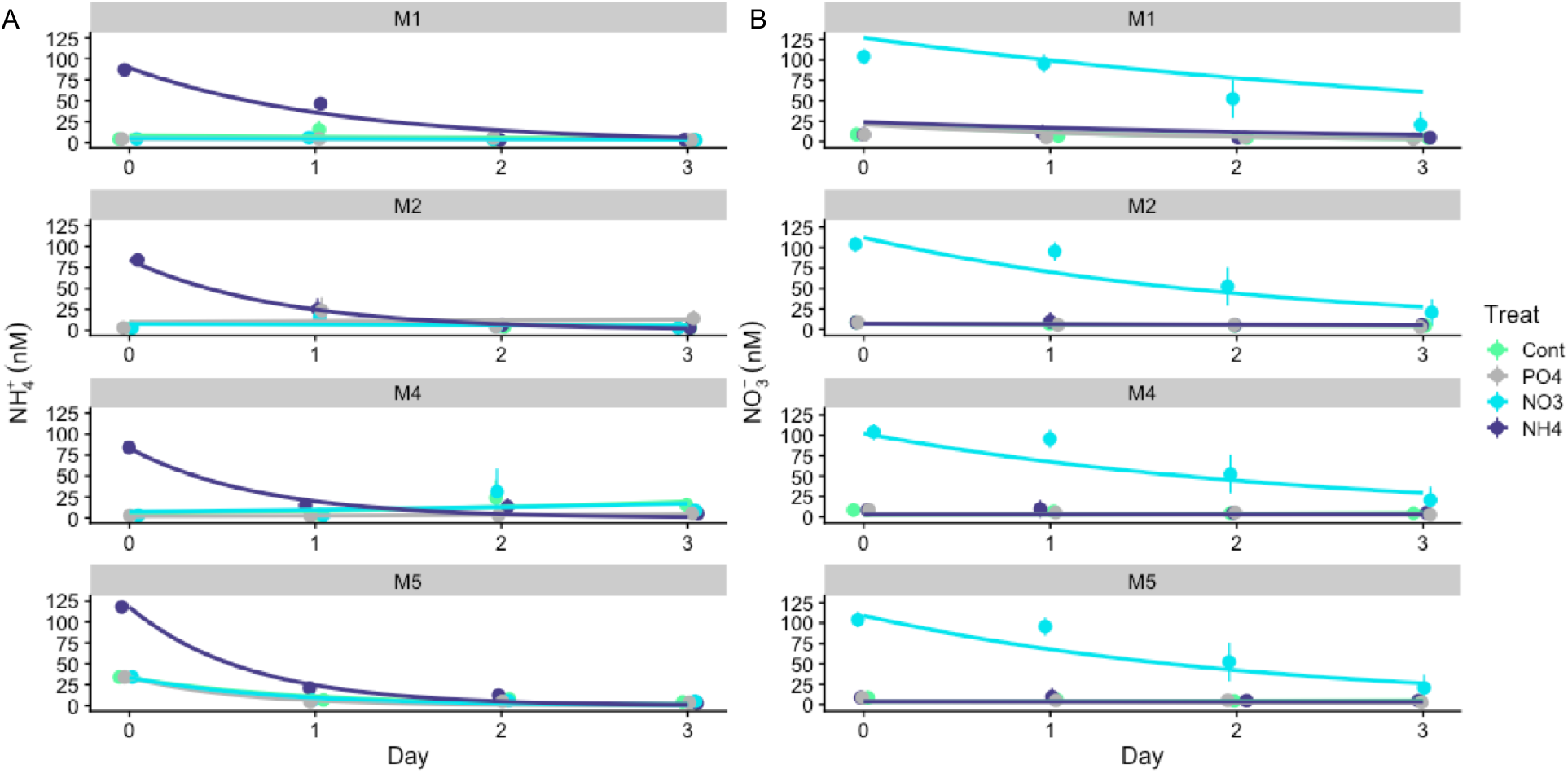
Temporal change in NH_4_^+^ and NO_3_^-^ concentrations of Ex. M1, M2, M4 and M5. (A) NH_4_^+^ concentration in the NH_4_^+^ treatment exponentially decreased during the experiment down to the detection limit of 6 nM on day 3. (B) NO_3_^-^ concentrations in the NO_3_^-^ treatment exponentially decreased during the experiment but enriched NO_3_^-^ was not always entirely consumed. Error bar shows a standard deviation of triplicate.

**Fig. S2.**
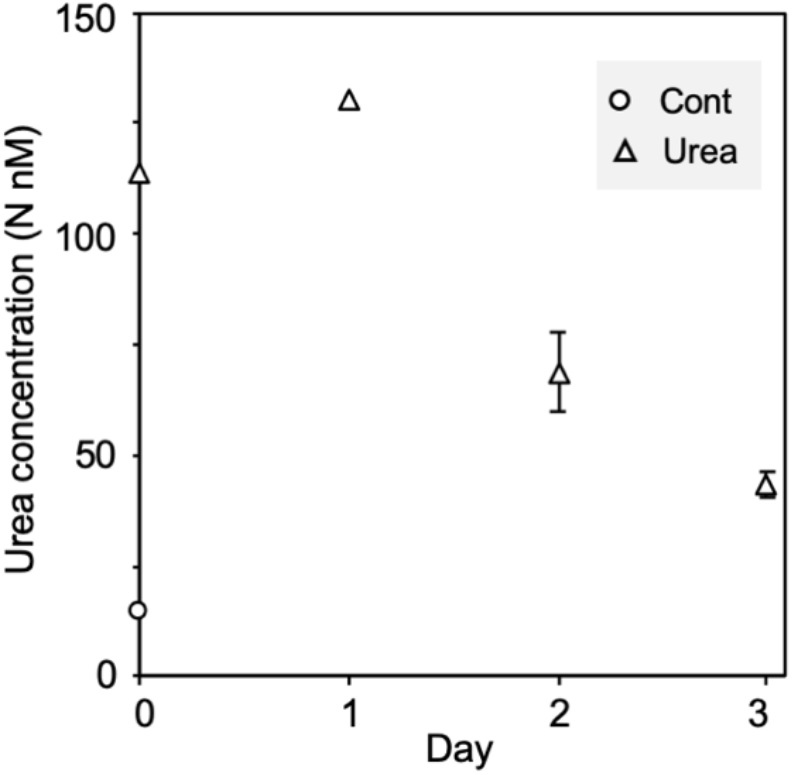
Temporal change in Urea-N concentration. Concentration in control was measured only at the initial. Error bar shows the standard deviation of triplicate samples.

**Fig. S3.**
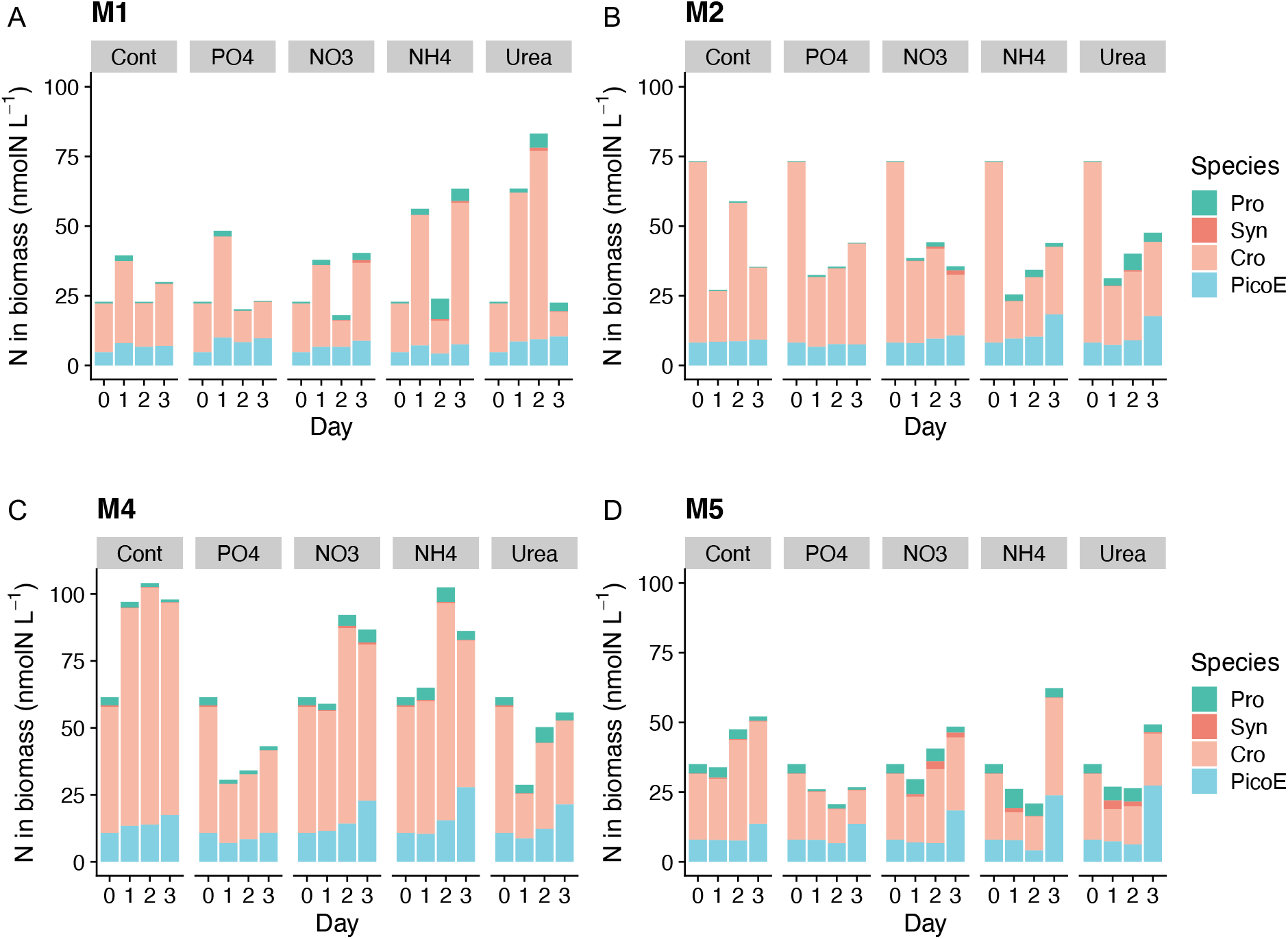
N in biomass in each treatment and its contribution of each phytoplankton group of experiment M1, M2, M4 and M5. Pro; *Prochlorococcus*, Syn; *Synechococcus*, Cro; *Crocosphaera*, PicoE; pico-eukaryotes.

**Fig. S4.**
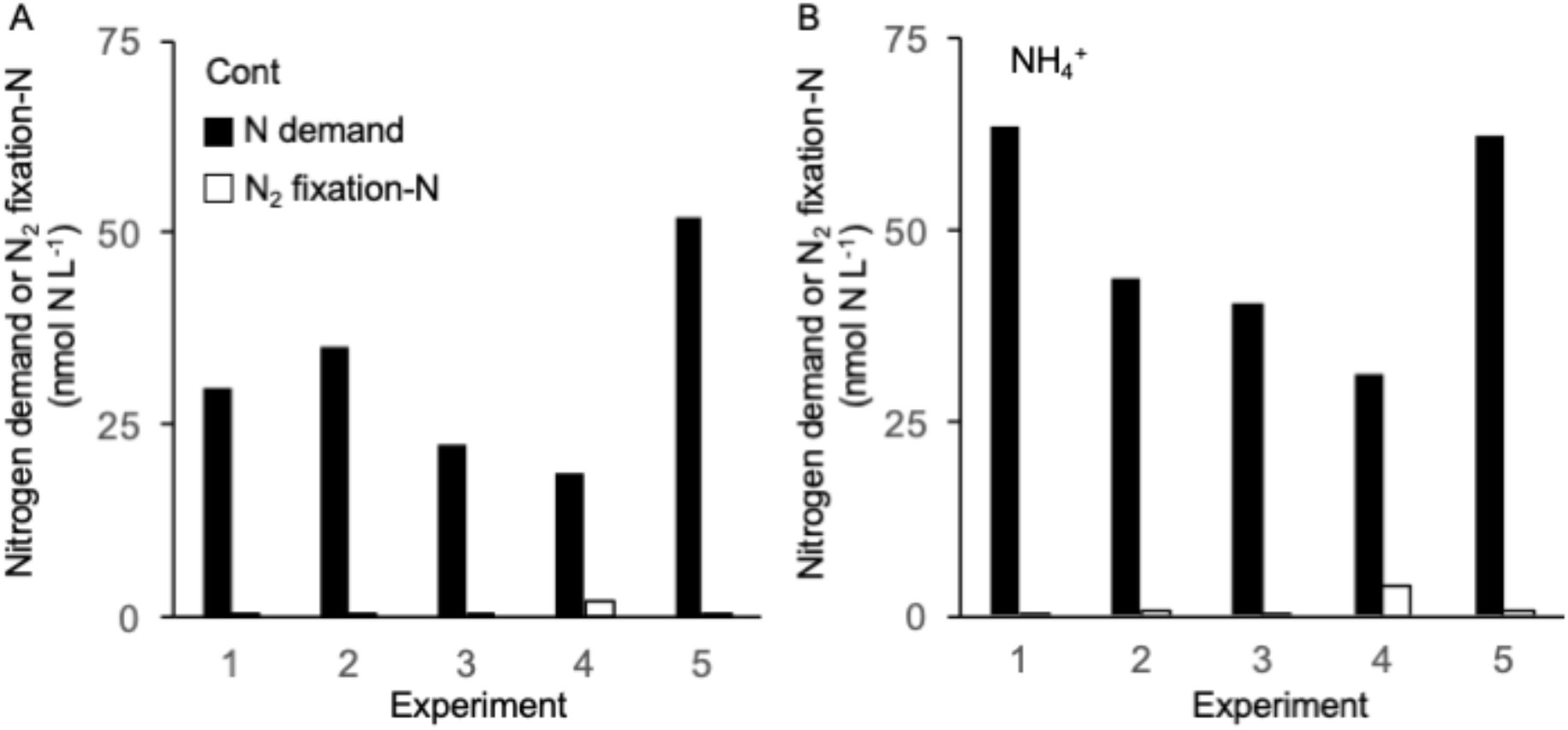
Nitrogen demand and N derived from N_2_ fixation in Control (A) and NH_4_^+^ treatment (B) for each experiment (M1-M5). Nitrogen demand is N in biomass in 3 days. N_2_ fixation rate was estimated from the reported maximum cellular N_2_ fixation rate 1.12 fmol N mol cell^- 1^ day^-1^ (valued obtained in day 3 in Fe + N treatment of Fe3 (Masuda et al Pre print)) and cell density.

**Fig. S5.**
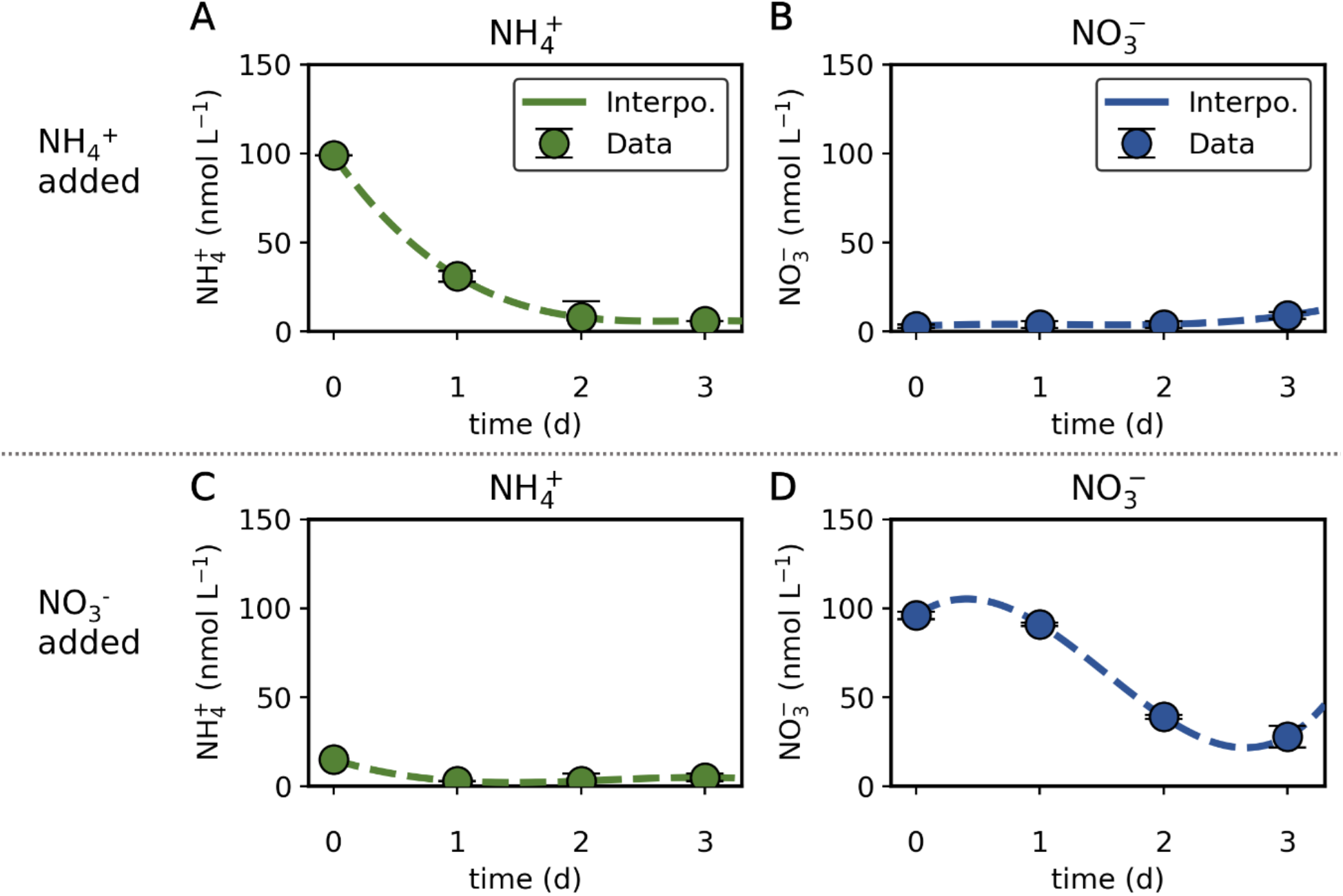
Measured NH_4_^+^ and NO_3_^-^ concentrations for NH_4_^+^ and NO_3_^-^ added cases. Dashed lines show quadratic interpolation. Data are from experiment M3.

**Fig. S6.**
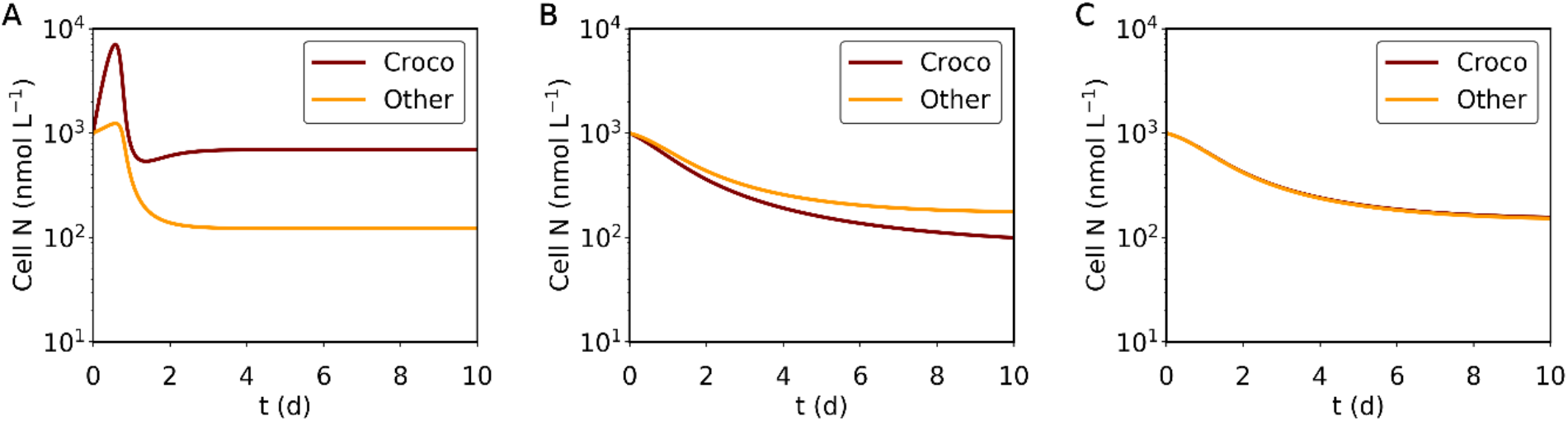
Simulated transition of cellular N in a simple ecosystem model for three different scenarios. (A) The concentrations for NH_4_^+^ and NO_3_^-^ are both 100 nmol L^-1^. (B)(C) The concentrations for NH_4_^+^ and NO_3_^-^ are both 1 nmol L^-1^. In only (C) *Crocosphaera* may acquire N via N_2_ fixation. Croco: *Crocosphaera*. Other: other phytoplankton. Parameters are based on NO_3_^-^ added case.

